# Differentiation of hypervirulent and classical *Klebsiella pneumoniae* with acquired drug resistance

**DOI:** 10.1101/2023.06.30.547231

**Authors:** Thomas A. Russo, Cassandra L. Alvarado, Connor J. Davies, Zachary J. Drayer, Ulrike Carlino-MacDonald, Alan Hutson, Ting L. Luo, Melissa J. Martin, Brendan W. Corey, Kara A. Moser, J. Kamile Rasheed, Alison L. Halpin, Patrick T. McGann, Francois Lebreton

## Abstract

Distinguishing hypervirulent (hvKp) from classical *Klebsiella pneumoniae* (cKp) strains is important for clinical care, surveillance, and research. Some combination of *iucA, iroB, peg-344, rmpA,* and *rmpA2* are most commonly used, but it is unclear what combination of genotypic or phenotypic markers (e.g. siderophore concentration, mucoviscosity) most accurately predicts the hypervirulent phenotype. Further, acquisition of antimicrobial resistance may affect virulence and confound identification. Therefore, 49 *K. pneumoniae* strains that possessed some combination of *iucA, iroB, peg-344, rmpA,* and *rmpA2* and had acquired resistance were assembled and categorized as hypervirulent hvKp (hvKp) (N=16) or cKp (N=33) via a murine infection model. Biomarker number, siderophore production, mucoviscosity, virulence plasmid’s Mash/Jaccard distances to the canonical pLVPK, and Kleborate virulence score were measured and evaluated to accurately differentiate these pathotypes. Both stepwise logistic regression and a CART model were used to determine which variable was most predictive of the strain cohorts. The biomarker count alone was the strongest predictor for both analyses. For logistic regression the area under the curve for biomarker count was 0.962 (P = 0.004). The CART model generated the classification rule that a biomarker count = 5 would classify the strain as hvKP, resulting in a sensitivity for predicting hvKP of 94% (15/16), a specificity of 94% (31/33), and an overall accuracy of 94% (46/49). Although a count of ≥ 4 was 100% (16/16) sensitive for predicting hvKP, the specificity and accuracy decreased to 76% (25/33) and 84% (41/49) respectively. These findings can be used to inform the identification of hvKp.

**Importance:** Hypervirulent *Klebsiella pneumoniae* (hvKp) is a concerning pathogen that can cause life-threatening infections in otherwise healthy individuals. Importantly, although strains of hvKp have been acquiring antimicrobial resistance, the effect on virulence is unclear. Therefore, it is of critical importance to determine whether a given antimicrobial resistant *K. pneumoniae* isolate is hypervirulent. This report determined which combination of genotypic and phenotypic markers could most accurately identify hvKp strains with acquired resistance. Both logistic regression and a machine-learning prediction model demonstrated that biomarker count alone was the strongest predictor. The presence of all 5 of the biomarkers *iucA, iroB, peg-344, rmpA,* and *rmpA2* was most accurate (94%); the presence of ≥ 4 of these biomarkers was most sensitive (100%). Accurately identifying hvKp is vital for surveillance and research, and the availability of biomarker data could alert the clinician that hvKp is a consideration, which in turn would assist in optimizing patient care.

## Introduction

Hypervirulent *Klebsiella pneumoniae* (hvKp) is capable of causing a variety of severe infectious syndromes in healthy hosts from the community as well as in hospitalized hosts (1, 2). By contrast, classical *K. pneumoniae* (cKp) primarily causes healthcare-associated infections (1). The ability to accurately identify hvKp has important clinical implications, such as identifying occult abscesses to enable source control and monitoring for endophthalmitis (3). Presently, clinical microbiology laboratories are unable to differentiate between these two pathotypes.

As a first step to be able to identify hvKp, our group previously designed a study in which two strain cohorts were assembled. Clinical criteria were used to assemble an hvKp-rich strain cohort; random blood isolates from geographical regions predicted to have a low prevalence of hvKp were used to create the cKp-rich strain cohort. Subsequent assessment of these strains identified 5 genotypic biomarkers that achieved a diagnostic accuracy of ≥0.95 for differentiating the hvKp-rich and cKp-rich strain cohorts, namely *iucA, iroB, peg-344, rmpA,* and *rmpA2.* This epidemiological analysis was experimentally validated in a murine sepsis model (4).

Subsequently, other investigators have used the presence of some of these markers to define a strain as being hypervirulent (5–8); often by the presence of some combination of *iucA* and *rmpA/A2* (9–12). However, it is important to note that *iucA, iroB, peg-344, rmpA,* and *rmpA2* biomarkers are linked on the canonical hvKp virulence plasmid pLVKP (13), which has been shown to be critical for the expression of the hvKp hypervirulent phenotype (14–16). The absence of some of these markers may reflect the presence of an incomplete virulence plasmid, which in turn could affect the hypervirulent phenotype. In our study (4) of the strains defined as hvKp (LD_50_ <10^7^ CFU) by virulence in the murine sepsis model, 80/85 (94%) strains possessed 5/5 of these biomarkers, 4/85 (5%) possessed 4/5 biomarkers (*rmpA2* was absent) and only 1/85 strains possessed 3/5 of these markers (*rmpA2*, *iucA* were absent) (4). Of note, of the strains that did not possess any of these 5 markers only 1/82 (1.2%) had an LD_50_ of <1 x 10^7^ CFU (the LD_50_ of that strain was 3.0 x 10^5^ CFU) (4). Further, the strains assessed in this initial study were identified by clinical syndrome irrespective of their antimicrobial susceptibility profile. As a result, these data did not adequately address the impact of possessing fewer than five of these markers or the effect of acquired antimicrobial resistance on the hypervirulent phenotype.

To address this knowledge gap, two collections of *K. pneumoniae* strains were assembled that possessed acquired antimicrobial resistance and the presence of the biomarkers *iucA, iroB, peg-344, rmpA,* and *rmpA2*. Forty-nine strains were identified that possessed 1-5 of these genes, their virulence was assessed in a CD1 murine subcutaneous challenge model, and these strains were categorized as hvKp or cKp (17). Next, we hypothesized that some combination of biomarker number, total siderophore production, mucoviscosity, Kleborate virulence score (18), and Mash/Jaccard distances (19, 20) to the canonical pLVPK could accurately determine the pathotype of these *K. pneumoniae* strains. Analysis by both stepwise logistic regression and a machine-learning based prediction model using a classification and regression tree (CART) demonstrated that the biomarker count alone proved to be the strongest predictor and was highly accurate. These data are of value for informing the development of a diagnostic test for the clinical microbiology laboratory as well as guiding surveillance, translational and clinical research, and patient care.

## Methods and Materials

### Bacterial strains

Forty-nine strains used in this study and selected genotypic and phenotypic characteristics are listed in Supplementary Table 1. Fifteen strains were obtained from the U.S. Centers for Disease Control and Prevention and 34 were obtained from the Walter Reed Army Institute of Research (WRAIR). All strains were stored at –70°C or –80°C prior to use.

### CD1 mouse subcutaneous (SQ) challenge infection models

Animal studies were reviewed and approved by the Veterans Administration Institutional Animal Care Committee and the University at Buffalo-SUNY and were carried out in strict accordance with the recommendations in the guidelines delineated in the “NIH Guide for the Care and Use of Laboratory Animals”(revised 1985) and the “Ethics of Animal Experimentation Statement” (Canadian Council on Animal Care, July 1980) as monitored by the Institutional Animal Care and Use Committee (IACUC). All efforts were made to minimize suffering. Veterinary care for the animals was supplied by the staff of the Veterans Administration Animal Facility under the direction of a fully licensed veterinarian. Studies were performed as described (4). The model is clinically relevant since hypervirulent *K. pneumoniae* (hvKp) may enter through micro-breaks in the skin, similar to *S. aureus*. Further, published data has established its ability to discriminate between cKp and hvKp strains (4). Lastly, we have established that this model produces similar results as intra-peritoneal challenge (17). The various challenge inocula and number of mice used for each strain are delineated in Supplementary Table S1. The animals were closely monitored for 14 days after challenge for the development of the study endpoints, survival or severe illness (*in extremis* state)/death, which was recorded as a dichotomous variable. Signs that were monitored and which resulted in immediate euthanasia using methods consistent with the recommendations of the American Veterinary Medical Association Guidelines included hunched posture, ruffled fur, labored breathing, reluctance to move, photophobia, and dehydration.

### Quantitative siderophore assay

Strains were grown individually overnight at 37°C in iron-chelated M9 minimal media containing casamino acids (c-M9-CA) (21) and culture supernatants were assessed using the chromeazurol S dye assay as described (22). A minimum of 3 biological assays with 3 technical repeats were performed and the results were reported as the mean ± the SD.

### Mucoviscosity assay

Since the hypermucoviscosity phenotype is more strongly associated with the hypervirulent phenotype compared to capsule (23, 24) this variable was chosen for study. Strains grown overnight in lysogeny broth (LB) were used to inoculate 10 mL of either LB or c-M9-CA plus added trace elements (5 µg/mL CaCl_2_, 1 µg/mL CoCl_2_, 20 µg/mL MgCl_2_, 10 µg/mL MnCl_2_) (c-M9-CA-te) to a starting OD_600_ of approximately 0.2. Strains were grown for 24 hours, and the assay was performed as described (17). A minimum of 3 biological assays were performed and the results were reported as the mean ± the SD.

### Growth/survival in human ascites fluid

Growth/survival experiments in human ascites were performed as previously described (25). The procedures for obtaining human ascites fluid were reviewed and approved by the Western New York Veterans Administration Institutional Review Board; informed consent for ascites fluid was waived because it was collected from de-identified patients who were undergoing therapeutic paracentesis for symptoms due to abdominal distension. These individuals were not being treated with antimicrobials and were not infected with human immunodeficiency virus, hepatitis B virus, or hepatitis C virus. The ascites fluid was cultured to confirm sterility, divided into aliquots, and stored at –80°C. For these experiments ascites obtained from a single individual was used.

### Sequencing and Assembly

All strains were sequenced from Illumina MiSeq or NextSeq platforms. CDC strains were assembled with Spades (Phoenix v1.) WRAIR strains were assembled with de novo assembly using Newbler (v2.7) with minimum thresholds for contig size and coverage set at 200Lbp and 49.5×, respectively. Long-read sequencing was completed on a subset of WRAIR isolates using with PacBio (Pacific Biosciences: Menlo Park, CA. USA) or MinION (Oxford Nanopore Technologies: Oxford, United Kingdom) platforms. Assembly was performed using the Trycycler tool (26). After filtering the bottom 5% of reads by quality score, reads were subsampled, with replacement, 12 times. Four assemblies were generated using Flye, Miniasm, and Raven using the 12 independent subset of reads. The consensus assembly was constructed with the assistance of Trycycler and was finally polished with short reads from Illumina MiSeq or NextSeq platforms.

### Identification of acquired antimicrobial resistance and virulence genes

Study strains possessed acquired antimicrobial resistance and some combination of the biomarkers *iucA, iroB, peg-344, rmpA,* and *rmpA2.* Kleborate (v2.3.2) was used to screen each isolate for any relevant virulence and antimicrobial resistance determinants (18). An agglomerative virulence score was determined by the presence or absence of the five genes: *iroB, iucA, peg-344*, and plasmid-borne *rmpA* and *rmpA2*. Possible loss-of-function mutations in the virulence biomarkers were predicted based on the Kleborate results as well as BLASTN and BLASTP analysis (Supplementary Table S1). All biomarkers except for *peg-344* are within the interrogative scope of Kleborate. The metabolite transporter *peg-344* was included in the agglomerative scoring due to its high association with fully virulent *K. pneumoniae* isolates (4). To calculate the prevalence of *iroB, iucA, peg-344*, *rmpA* and *rmpA2* in MDR *K. pneumoniae,* an unbiased collection of 3,123 clinical isolates (27) from global origins was queried using BLASTN and Kleborate (Supplementary Table S2). Strains also underwent antimicrobial susceptibility testing using reference broth microdilution with panels that were prepared in-house according to the Clinical and Laboratory Standards Institute (CLSI) guidelines (28) and stored at –70°C until use. Results of testing were interpreted according to guidance provided in CLSI document M100 (29), with the exception of tigecycline, for which criteria established by the U.S. Food and Drug Administration were applied. *E. coli* ATCC 25922, *P. aeruginosa* ATCC 27853, and *K. pneumoniae* ATCC 700603 were used for quality control.

### Mash and Jaccard distance indices

To determine genetic relatedness of each isolates’ virulence plasmid to the canonical pLVPK virulence plasmid (13), Mash distance (19) and Jaccard similarity indices (20) were calculated between closed plasmid sequences containing virulence biomarkers and pLVPK for the 29 closed genomes (hvKp N=9, cKp N=20). Mash distance calculation was performed with the dependency-free binary provided freely by the developers (https://github.com/marbl/mash). The Jaccard similarity is defined by the proportion of genes shared by two comparator plasmids over the set of all observed genes in the two plasmids. Hence, the Jaccard distance is the complement of the similarity value and represents the proportion of all observed genes that are not shared in the two comparator plasmids. The Jaccard metric was calculated from Roary output (30), which provided a gene presence or absence table from a set of annotated input genome assemblies. The Vegan package in R was used to calculate Jaccard distance between each pairwise comparison of virulence plasmids to pLVPK.

### Comparative genomics

Two strains (AKp14 and AKp35) carried all five biomarkers and did no exhibit an hypervirulent phenotype in the murine model. Gene content analysis was performed using Roary (30) to investigate possible genetic factors involved in the loss of virulence. For strain AKp14 (ST23), we compared its genome to those of all ST23 confirmed hvkp in this study (AKp4, AKp10, AKp25, AKp27 and RCA1) (Supplementary Table S3). For strain AKp35 (ST412), there was no other ST412 with available virulence data to compare. Instead, its genome was compared to all the 16 strains (various ST) with confirmed hvKp phenotype in this study (Supplementary Table S3).

### Statistical Analyses

The LD_50_ was estimated using a logistic regression model with the factors for strain and inoculum (CFU/mL). The estimated LD_50_ was given as 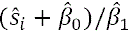 where 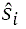 is the strain effect, 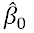 is the intercept and 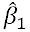 is the regression parameter associated with inoculum. A model fit statistical analysis was performed (c-stat). C was equal to 0.879 (C = 1.0 is a perfect fitting model), which supports that the model was a good fit. The k-means cluster analysis was employed to examine whether distinct groupings could be formed based on the LD_50_ values, using two, three, or more clusters. The analysis revealed two well-defined clusters that were categorized as cKP (LD_50_ of > 1 x 10^7^ CFU) and hvKP (LD_50_ of ≤ 1 x 10^7^ CFU).

Mucoviscosity when grown in LB or c-M9-CA-te, biomarker numbers (*iucA*, *iroB*, *peg-344*, *rmpA*, *rmpA2*), Mash and Jaccard distances, and Kleborate scores for the hvKp and cKp strain cohorts were initially compared using a Mann-Whitney test. Siderophore production for the two cohorts was compared utilizing an unpaired t test (Prism version 9.5.1).

Exact logistic regression was used to model the association of class with each marker of virulence and to estimate the area under the ROC curve (AUC) in both a univariate and multivariate fashion. The AUC ranges from 0.5 to 1 and is a measure of the percent agreement between the observed and predicted values. A value of AUC=0.5 implies no predictive utility. A value of 1.0 implies 100% accuracy in terms of prediction (SAS).

A machine-learning based prediction model using a classification and regression tree (CART) was used to examine the prediction of the strain cohorts cKP (LD_50_ of > 1 x 10^7^ CFU) and hvKP (LD_50_ of ≤ 1 x 10^7^ CFU) as a function of count (sum of positive *iucA*, *iroB*, *peg-344*, *rmpA* and *rmpA2* markers), quantitative siderophore production, mucoviscosity when grown in LB or c-M9-CA-te, Mash Distance, Jaccard Distance, and Kleborate Score. The method of cost-complexity pruning was utilized to determine the final tree fit (SAS).

## Results

### Strain cohorts

Forty-nine *K. pneumoniae* strains were collected from 8 countries (n= 30 from USA; n= 4 from Germany; n= 3 from Ukraine; n= 3 from Middle East; n= 2 from Thailand; n= 5 from Afghanistan; n= 1 from Uganda; n= 1 from Turkey). Twenty-eight sequence types that possessed some combination of the biomarkers *iucA, iroB, peg-344, rmpA,* and *rmpA2* and antimicrobial resistance other than the ubiquitous ampicillin resistance were chosen for study (Supplementary Table S1). These included the ST23, ST11, ST29, and ST268 hvKp lineages, but also the “high-risk” emerging multi-drug resistant (MDR) lineages ST395, ST307, and ST15. Of these 49 strains 34 (69%) possessed an extended-spectrum ß-lactamase (ESBL) and 20 (41%) possessed a carbapenemase; the mean number of resistance genes was 7.51 (range 1-14) (Supplementary Table S1). Of these 49 strains, 17 (35%) possessed all 5 biomarkers, 7 (14%) possessed 4 biomarkers, 19 (39%) possessed 2 biomarkers, and 6 (12%) possessed 1 biomarker (Supplementary Table S1). To place this in perspective, the natural prevalence of biomarkers in an unbiased collection of 3,123 MDR isolates (27) from global origins were assessed. The most represented biomarkers were *peg-344* (8% of isolates), *iucA* (5%) *rmpA2* and *rmpA* (each at 4%) and *iroB* (1%). But only 0.9% (n= 28), 2.7% (n= 84), 1% (n= 30) and 4% (n= 127) of isolates carried 5, 4, 3 or ≤ 2 biomarkers respectively (Supplementary Table S2).

Next, these 49 strains underwent pathotype characterization as defined by virulence in the outbred CD1 SQ challenge model. hvKp were defined by an LD_50_ of ≤1 x 10^7^ CFU and cKp was defined by an LD_50_ of > 1 x 10^7^ CFU (17). Sixteen (33%) strains were categorized as hvKp (mean log LD_50_ 4.06 ± 1.91) and 33 (67%) as cKp (mean log LD_50_ 8.91 ± 0.31) (Figure 1A, Table 1).

**Figure 1.**
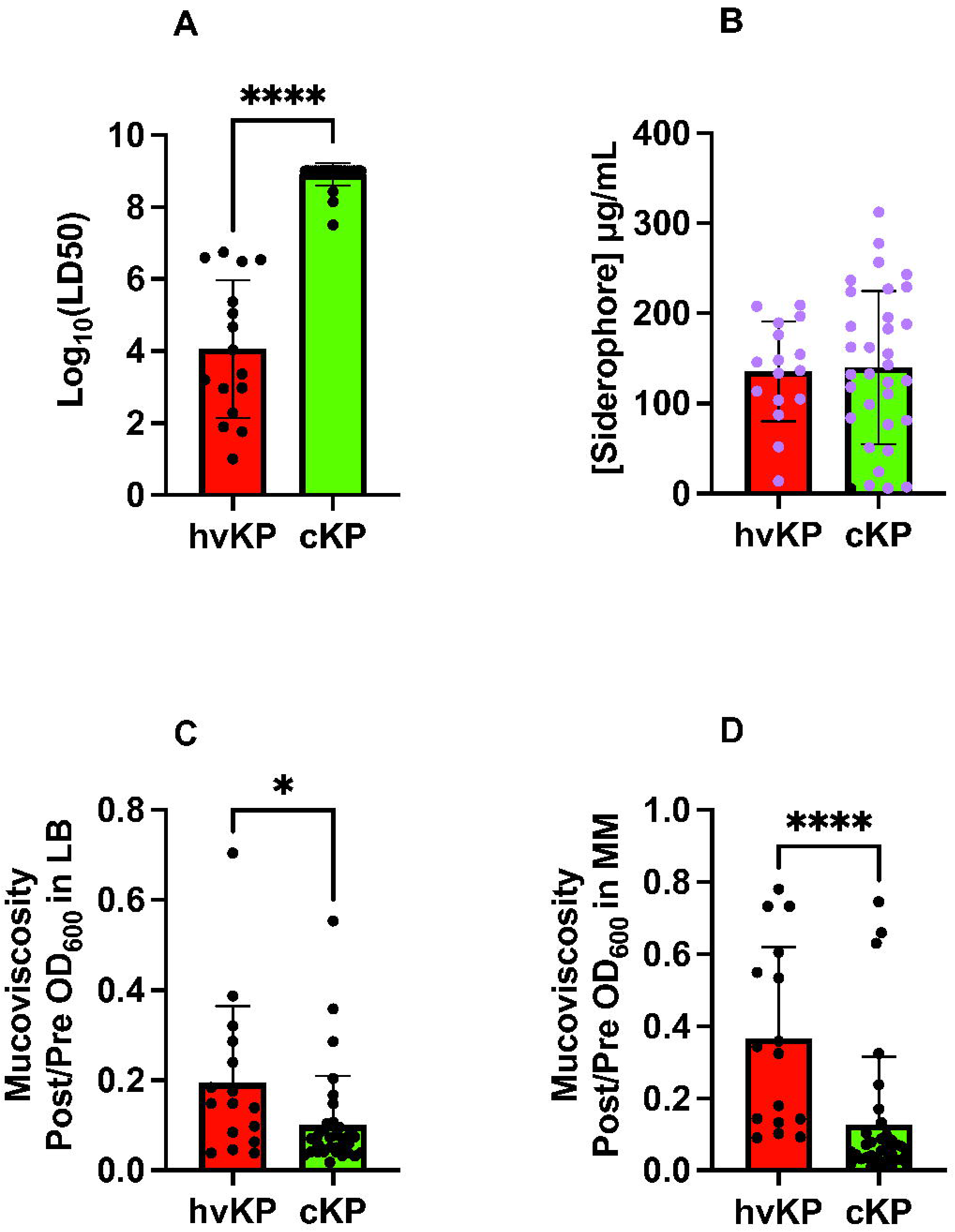
Phenotypic markers. Forty-nine strains comprising the hvKp (n=16) and cKp (n=33) cohorts underwent assessment for LD_50_ in the outbred CD1 murine subcutaneous challenge infection model (Panel A), *in vitro* quantitative siderophore production when grown in c-M9-CA, black symbols represent strains that do not possess *iucA* (Panel B), and *in vitro* quantitative mucoviscosity when grown in either LB (Panel C), or c-M9-CA-te medium (Panel D). For Panel A see Supplementary Table S1 for challenge inocula and animal number for each strain. For Panels B a minimum of three biologic with two technical repeats was performed for each strain. For Panels C and D a minimum of three biologic repeats was performed for each strain. Strain cohorts were compared via Mann-Whitney except siderophore production was compared by an unpaired t test (Prism version 9.5.1). <u>Panel A</u>, **** P < 0.0001. <u>Panel C</u>, * P = 0.0113, <u>Panel D</u>, **** P < 0.0001

**Table 1.**
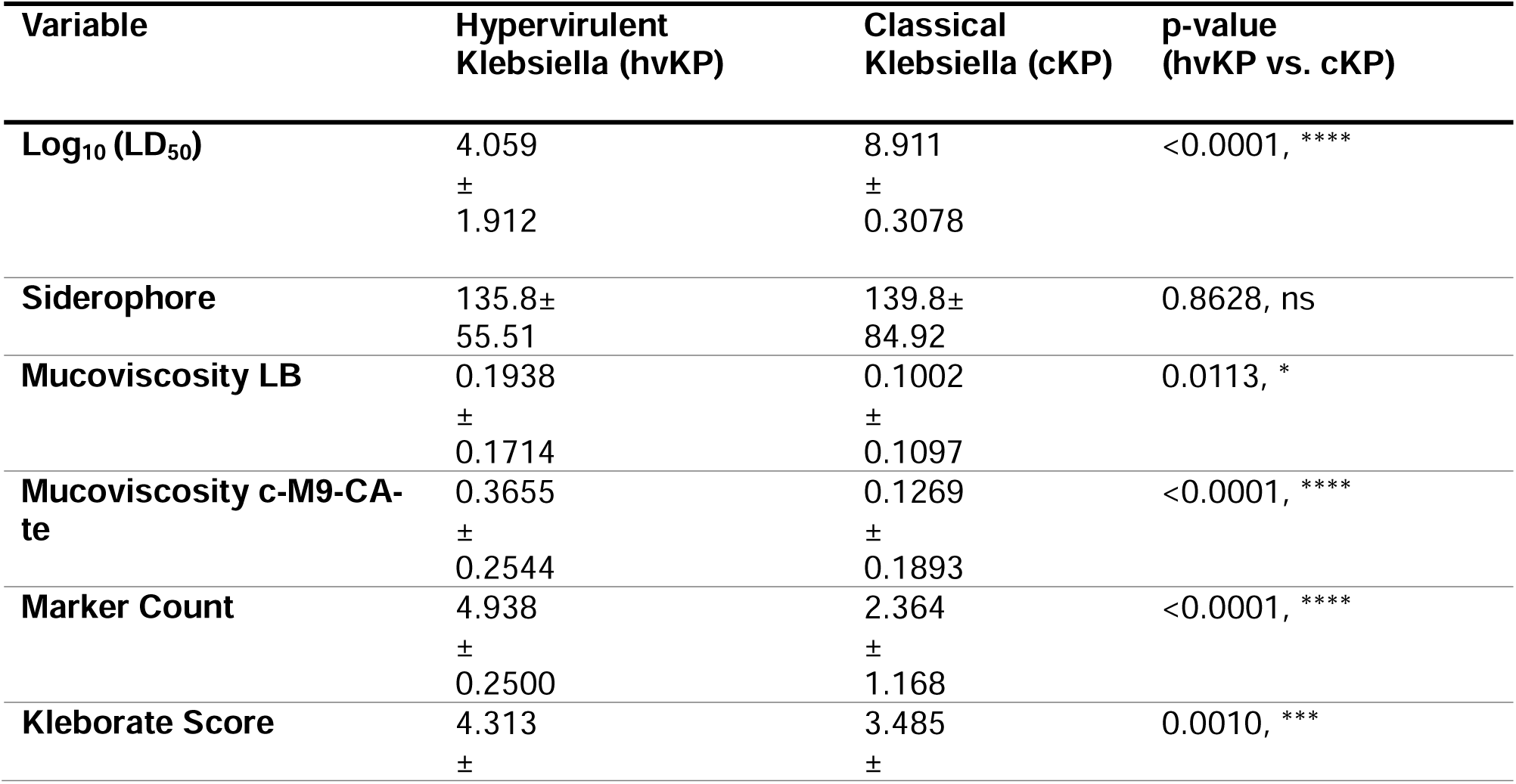
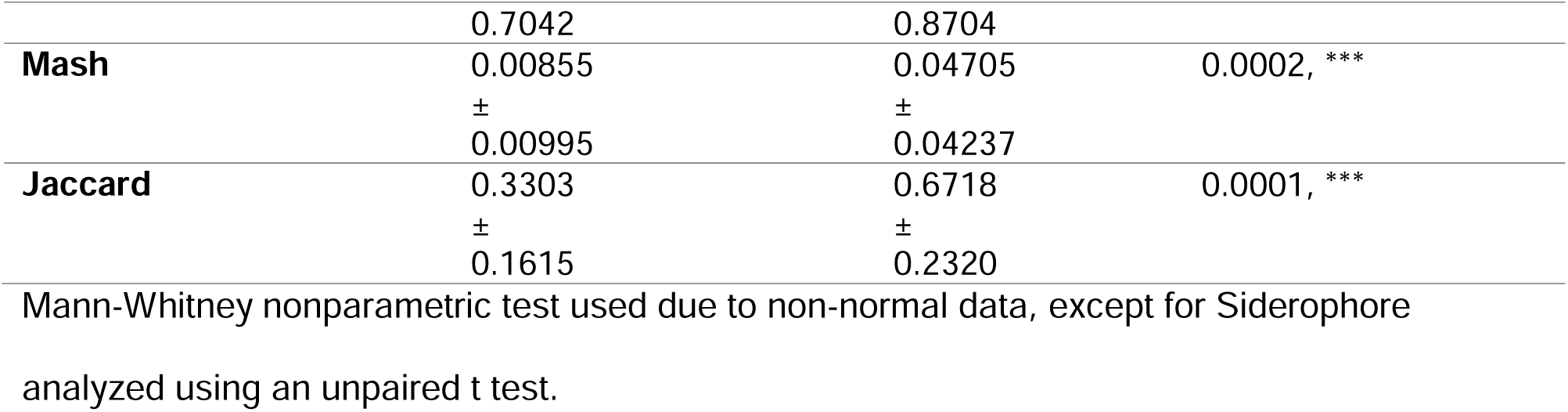
Values for genotypic and phenotypic markers.

### Phenotypic assessment of the hvKp and cKp strain cohorts

Previous studies have demonstrated that quantitative siderophore production and mucoviscosity assessment held promise for differentiating hvKp from cKp strains (4, 17). Therefore, we evaluated these phenotypes for all 49 strains in the hvKp and cKp strain cohorts.

Quantitative siderophore production was similar for the hvKp (135.8 ± 55.5 µg/mL) and cKp (139.8 ± 84.9 µg/mL) strain cohorts (Figure 1B, Table 1). When grown in LB medium, the hvKp strain cohort (0.194 ± 0.171) was significantly more mucoviscous than the cKp strain cohort (0.1002 ± 0.1097) (P = 0.0113) (Figure 1C, Table 1). When grown in c-M9-CA-te medium, the hvKp (0.366 ± 0.254) was significantly more mucoviscous than the cKp strain cohort (0.1269 ± 0.1893) (P < 0.0001) (Figure 1D, Table 1). These data demonstrate that quantitative siderophore production was unable to distinguish the hvKp and cKp strain cohorts, but quantitative mucoviscosity could.

### Genotypic Assessment of the hvKp and cKp strain cohorts

Previous studies demonstrated that the presence of *iucA*, *iroB*, *peg-344*, *rmpA*, and *rmpA2*, which are linked on the hvKp-specific virulence plasmid possessed a >95% accuracy for differentiating the hvKp-rich and cKp-rich strain cohorts (4). Therefore, all 49 strains in the hvKp and cKp strain cohorts were assessed for these genes and their Kleborate virulence score was calculated.

The hvKp strain cohort has a significantly higher marker count (some combination of *iucA*, *iroB*, *peg-344*, *rmpA*, and *rmpA2*) (4.94 ± 0.25) compared to the cKp strain cohort (2.36 ± 1.17) (P < 0.0001) (Figure 2A, Table 1). Of note, all 25 strains that possessed only one or two markers were categorized as cKp based on the murine infection model (Figure 3A, Supplementary Table S1) (31). Of these, 19/25 (76%) possessed *iucA* and *rmpA2* and 5/25 (20%) possessed *iucA* alone. In addition, the Kleborate virulence score was assessed, which is defined as the genes for aerobactin (3 points), yersiniabactin (1 point), and colibactin (1 point). The average Kleborate score was significantly different for the hvKp (4.3 ± 0.70) compared to the cKp (3.5 ± 0.87) (P= 0.001) strain cohort (Figure 2D, Table 1). Individually, within the 16 hvKp strains, 14 (87%) received a Kleborate score of five or four, and the remaining two scored as three on the virulence scale (Figure 3B, Supplementary Table S1). However, a virulence score of five or four also represented 50% of cKp isolates (17/33) (Figure 3B, Supplementary Table S1).

**Figure 2.**
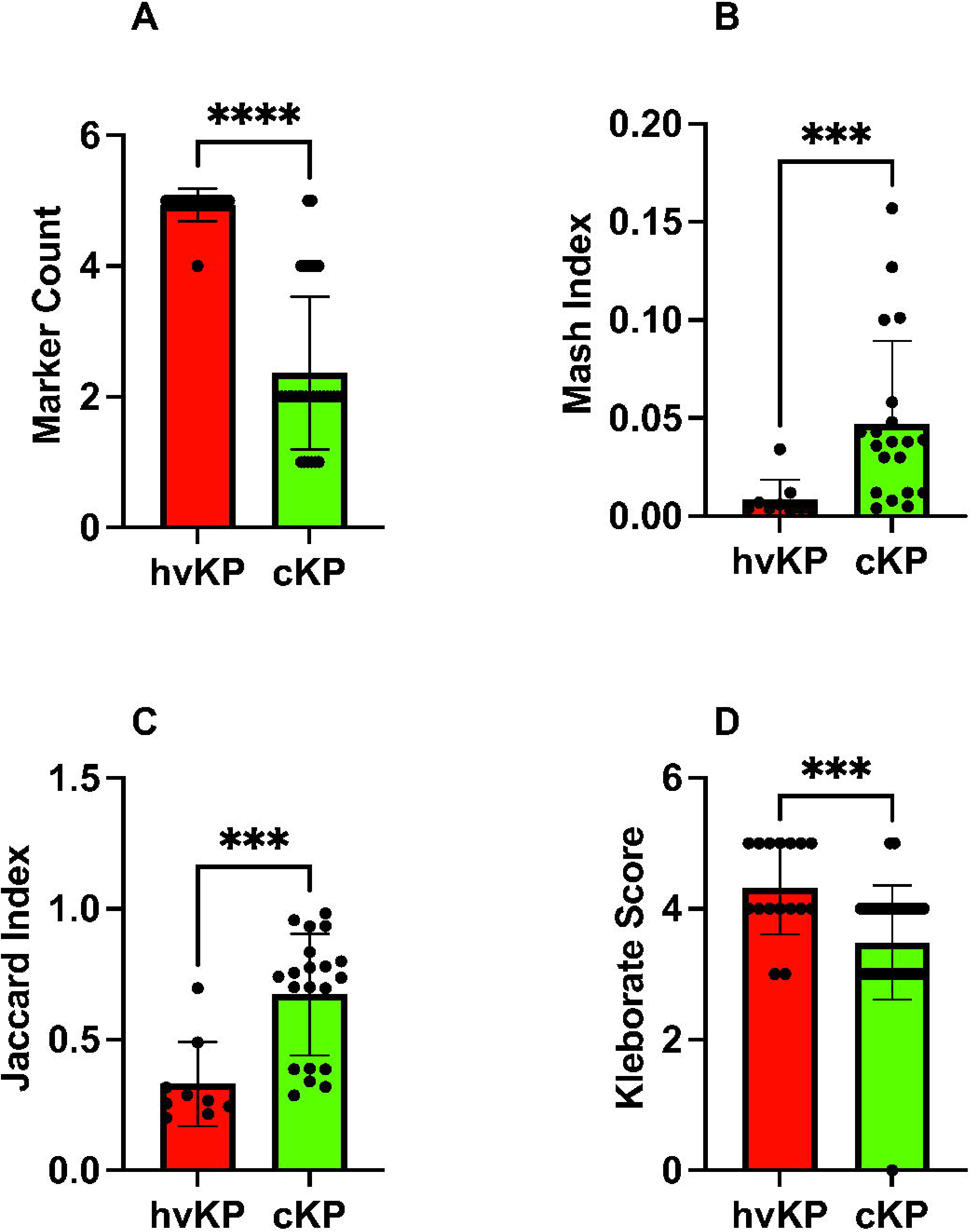
Genotypic markers. Forty-nine strains comprising the hvKp (n=16) and cKp (n=33) cohorts underwent assessment for marker count (Panel A), 29 strains comprising the hvKp (n=9) and cKp (n=20) cohorts underwent assessment for Mash distance (Panel B), and for Jaccard distance (Panel C), and all 49 strains underwent assessment for Kleborate score (Panel D). Strain cohorts were compared via Mann-Whitney (Prism version 9.5.1). <u>Panel A</u>, **** P = <0.0001, <u>Panel B</u>, *** P = 0.0002, <u>Panel C</u>, *** P = 0.0001, Panel D, *** P = 0.001

**Figure 3.**
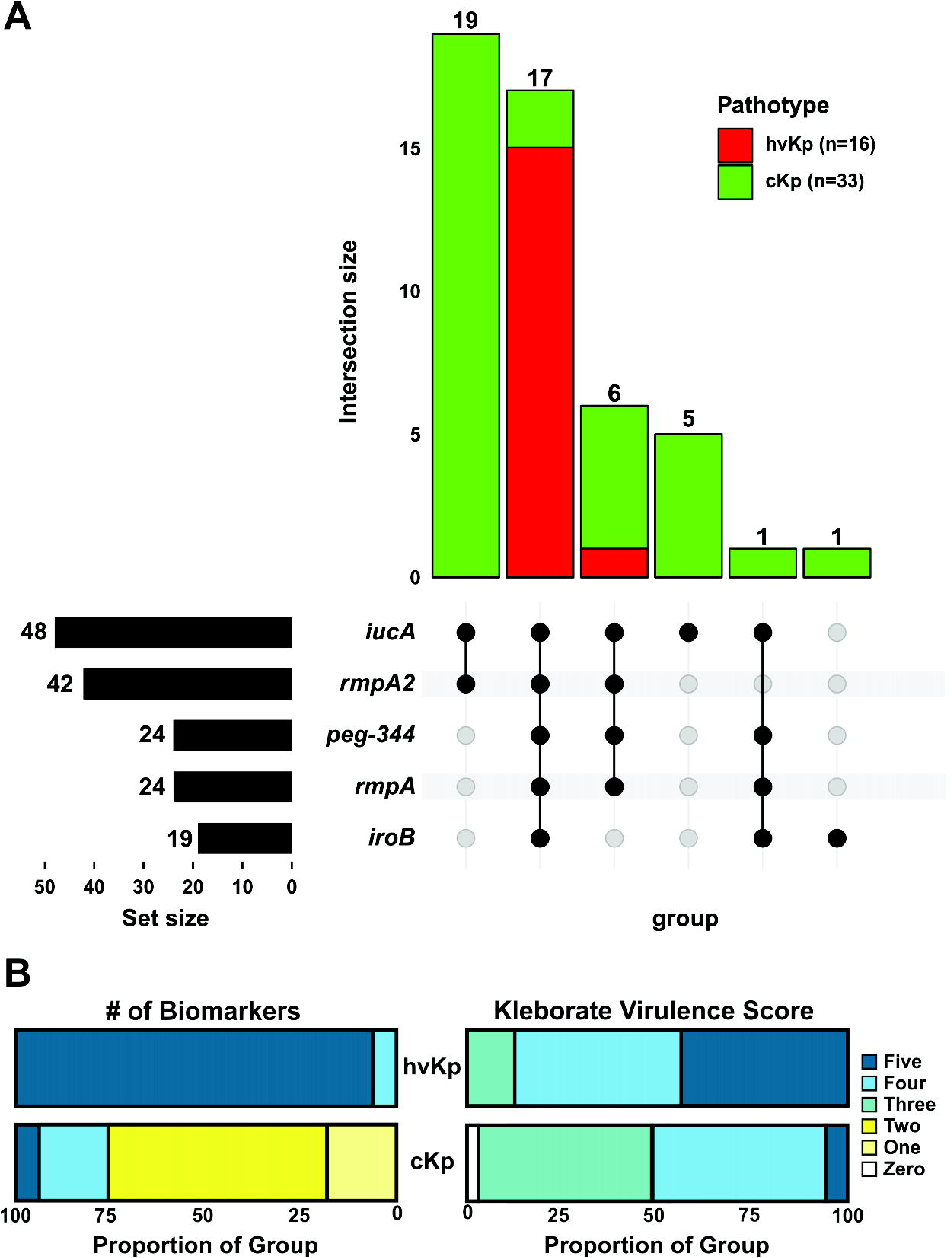
Biomarker distribution. <u>Panel A</u>. UpSet graph for the 49 strains comprising hvKp (n=16) and cKp (n=33) was generated to facilitate the visualization of the presence or absence of the biomarkers *iucA*, *iroB*, *peg-344*, *rmpA*, and *rmpA2*. Black circles designate the presence of a given marker whereas grey circles designate its absence. Pathotypes are color coded. The number above each bar represents the number of strains that possess that marker configuration. <u>Panel B</u>: Results are shown as the distribution in proportions of each biomarker count (1–5) and Kleborate virulence score (0–5) for the hvKp and cKp cohorts. The biomarkers count is the presence of some combination of *iucA*, *iroB*, *peg-344*, *rmpA*, and *rmpA2.* The Kleborate virulence score is calculated by the presence of the genes that encode aerobactin (3 points), yersiniabactin (1 point), and colibactin (1 point).

Besides the presence of a biomarker gene, possible loss-of-function mutations were also characterized (Supplementary Table S1). Compared to reference allele, 88% (37/42) of the isolates with *rmpA2* carried a predicted truncated allele. This was observed at similar frequencies in cKp (88% 23/26) and hvKp (87% 14/16) isolates with identical late frameshifts (E96fs*101, E96fs*115 and N118fs*127) being the most represented. Similarly, predicted truncations in *rmpA* were observed in 27% (6/22) of isolates carrying this biomarker despite all being categorized as hvKp phenotypically. For other biomarkers, an early stop codon (K7*) was identified in *iucA* in 5 cKP strains and a single strain carried a late frameshift (T291fs*298) in *peg-344* but was still categorized as hvKp based on the mouse survival data.

Beyond individual biomarkers, it was hypothesized that carriage of an hvKp virulence plasmid would also be predictive of the hvKp pathotype as measured by Mash/Jaccard distances compared to the canonical virulence plasmid pLVKP. Mash/Jaccard distances for pLVLP-like virulence plasmids from the 29 closed genomes (hvKp N=9, cKp N=20) were measured. The Mash distance was significantly different for the hvKp (0.0086 ± 0.001) compared to cKp (0.04705 ± 0.042) strain cohort (P = 0.0002) (Figure 2B, Table 1). Likewise, the Jaccard distance was significantly different for the hvKp (0.33 ± 0.16) compared to the cKp (0.6718 ± 0.2320) (P = 0.0001) (Figure 2C, Table 1). Each pLVPK-like plasmid can be compared to the canonical pLVPK sequence by its Mash and Jaccard distance. The two distances are highly correlated, but do not scale linearly (Figure 4). Figure 4 show scatterplots constructed using these two distances whereby each point (n=29) represents a strain’s pLVPK-like plasmid. As Mash and Jaccard distances approach 0, (i.e., the strain’s pLVPK-like sequence and shared gene content is more similar to pLVPK), the strain is more likely to belong to a hvKp pathotype (Figure 4, top left panel, Pathotype). Viewing from different perspectives, we observe that pLVPK-like plasmids that are more similar to pLVPK, are also more likely to carry all five biomarkers within that plasmid (Figure 4 top right panel, # of Biomarkers). Within the same scope, we can extend generalizations at the genome level from the sequences of their respective pLVPK-like plasmids. In general, strains that carry a pLVPK-like that is highly similar to pLVPK are more likely to score ≥3 in Kleborate virulence (Figure 4 bottom left panel, Kleborate Virulence Score) and are also less likely to carry ESBLs/carbapenemases (Figure 4 bottom right panel, AMR) elsewhere in the genome.

**Figure 4.**
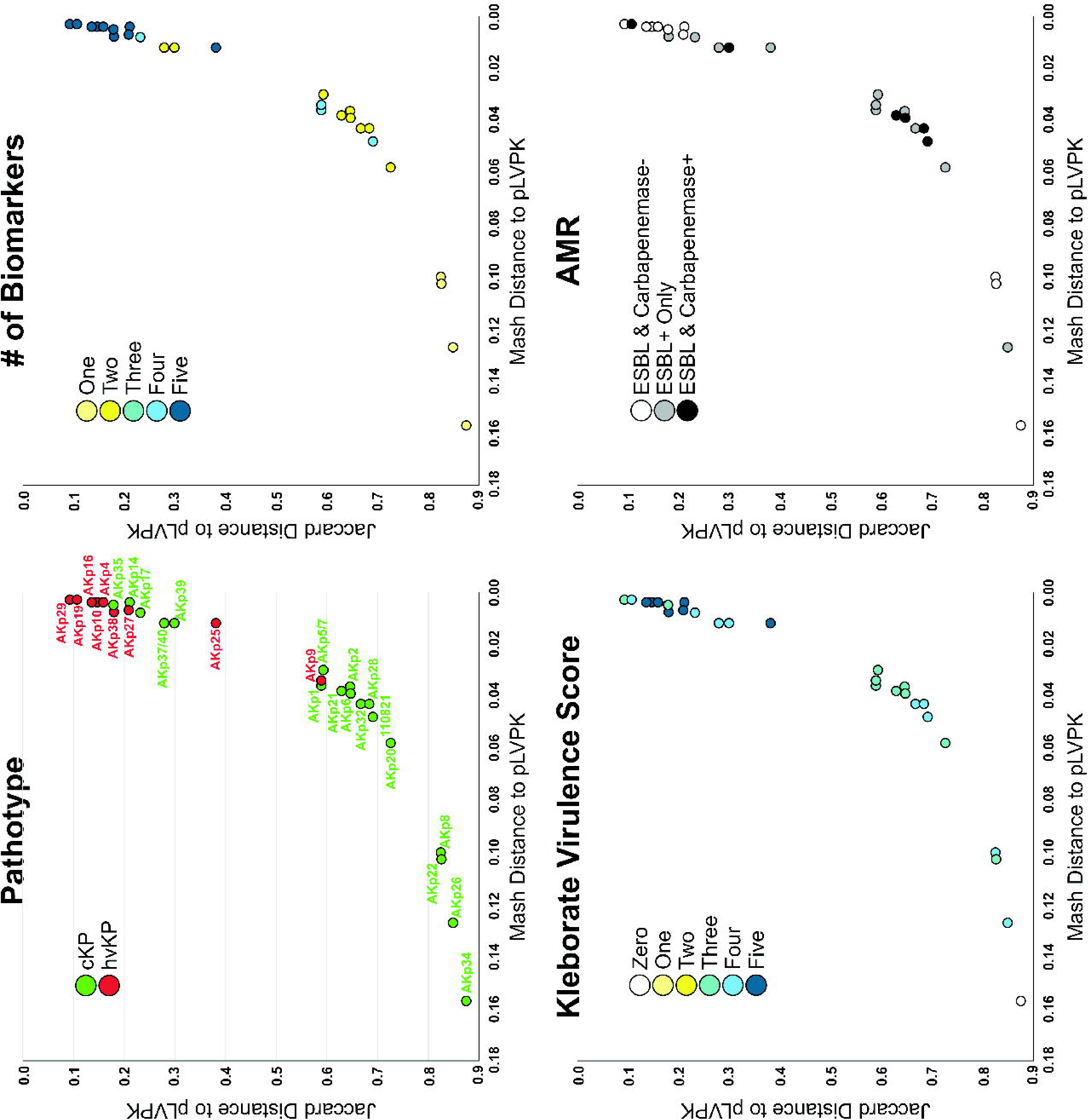
Mash and Jaccard Distances. The pLVKP-like virulence plasmid of 29 strains comprising the hvKp (n=9) and cKp (n=20) cohorts, for which long-read sequencing was obtained thereby enabling a closed genome/plasmid, was compared to the canonical pLVPK sequence. Scatterplots of Mash/Jaccard distance to pLVPK were plotted for each strain and colored by pathotype, number of biomarkers (some combination of *iucA*, *iroB*, *peg-344*, *rmpA*, *rmpA2*), Kleborate virulence score (presence of genes that encode aerobactin (3 points), yersiniabactin (1 point), and colibactin (1 point)), and antimicrobial resistance (AMR, presence of ESBL/carbapenemase genes).

### Logistic regression and CART prediction models

Biomarker number, total siderophore production, mucoviscosity, Kleborate virulence score, and Mash/Jaccard distances to the canonical pLVPK were analyzed by both stepwise logistic regression and a CART model to determine which factor or combination of factors were most accurate for predicting the hvKp and cKp strain cohorts. The strongest predictor in both models was the presence of all five of the biomarkers *iucA*, *iroB*, *peg-344*, *rmpA*, and *rmpA2* (Table 2). In the logistic regression model the AUC for biomarker count was 0.962 (P = 0.004). In that model the Jaccard distance was the next most predictive with an AUC of 0.919 (P < 0.001). Likewise, the CART model predicted biomarker count alone as most accurate. Using a cut-point of possession of all 5 biomarkers resulted in a sensitivity for predicting hvKP of 94% (15/16), a specificity of 94% (31/33), and an overall accuracy of 94% (46/49). A count of ≥ 4 resulted in a 100% (16/16) sensitivity for predicting hvKp, but specificity was decreased to 76% (25/33) and accuracy decreased to 84% (41/49). The Jaccard distance, Mash distance, mucoviscosity (c-M9-CA-te), Kleborate score, mucoviscosity (LB), and siderophore concentration demonstrated accuracies of 90%, 86%, 80%, 78%, 71%, and 49% respectively (Table 2).

**Table 2.**
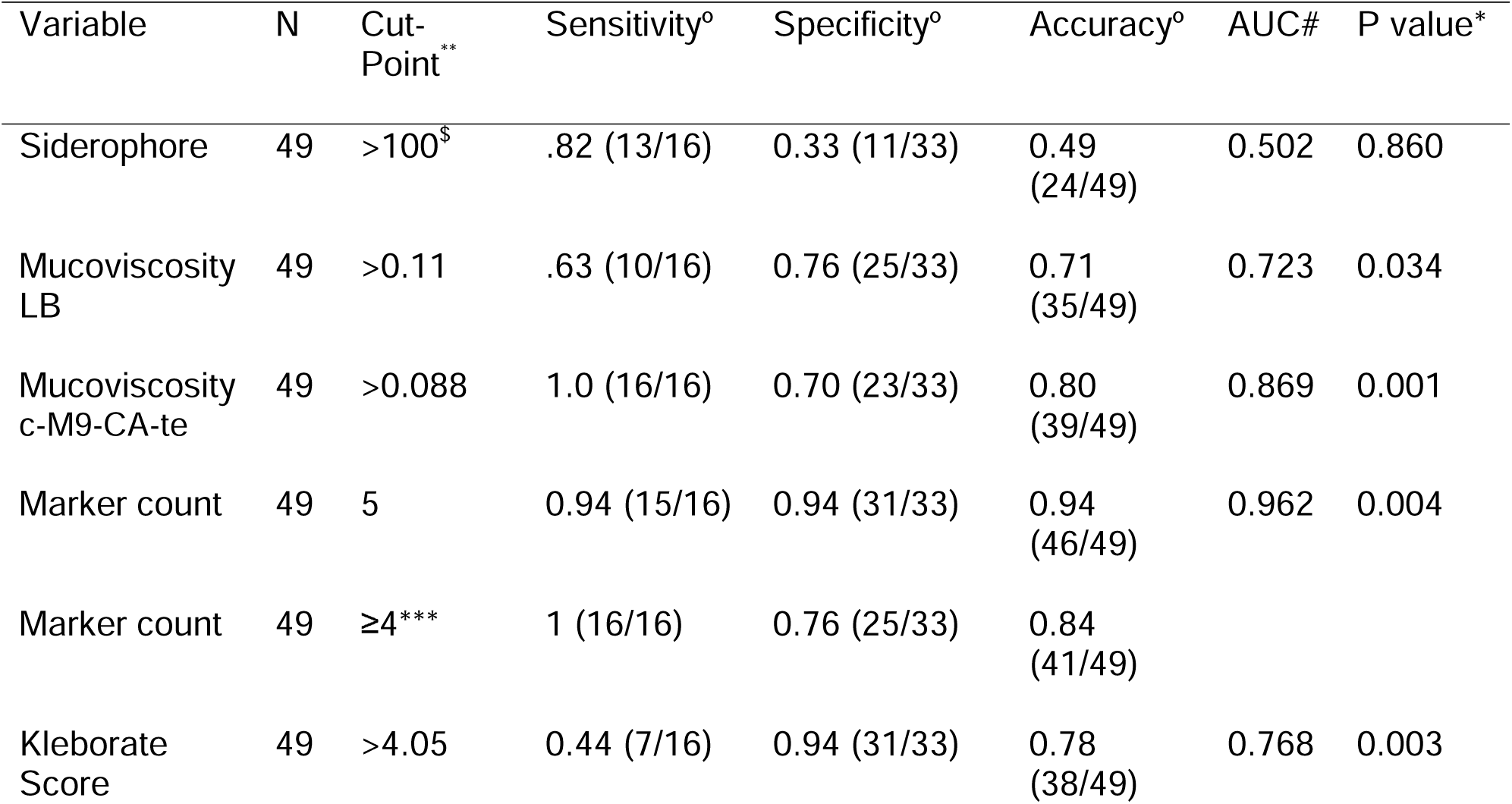
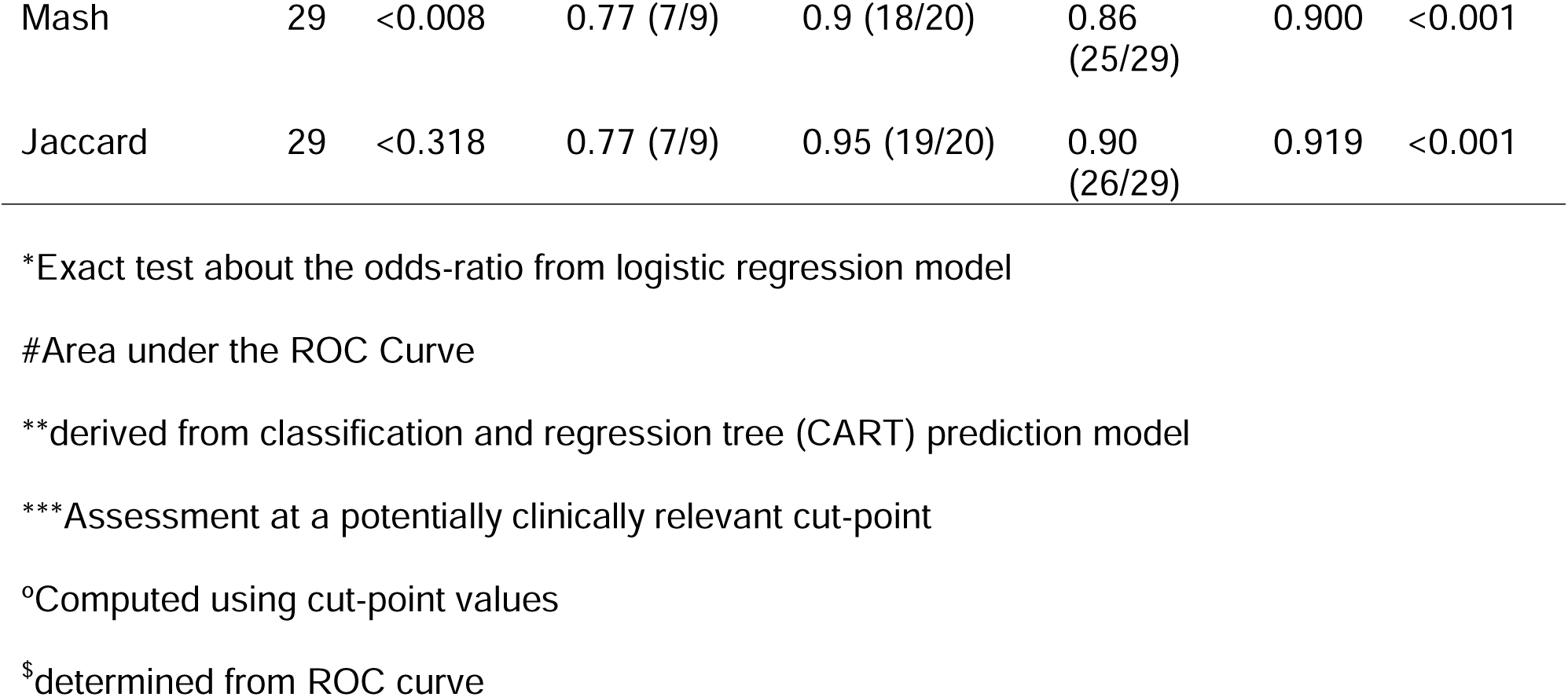
Logistic regression and CART prediction models.

| Variable | N | Cut-Point** | Sensitivity<sup>o</sup> | Specificity<sup>o</sup> | Accuracy<sup>o</sup> | AUC# | P value* |
| --- | --- | --- | --- | --- | --- | --- | --- |
| Siderophore | 49 | >100<sup>\$</sup> | .82 (13/16) | 0.33 (11/33) | 0.49 (24/49) | 0.502 | 0.860 |
| Mucoviscosity LB | 49 | >0.11 | .63 (10/16) | 0.76 (25/33) | 0.71 (35/49) | 0.723 | 0.034 |
| Mucoviscosity c-M9-CA-te | 49 | >0.088 | 1.0 (16/16) | 0.70 (23/33) | 0.80 (39/49) | 0.869 | 0.001 |
| Marker count | 49 | 5 | 0.94 (15/16) | 0.94 (31/33) | 0.94 (46/49) | 0.962 | 0.004 |
| Marker count | 49 | ≥4*** | 1 (16/16) | 0.76 (25/33) | 0.84 (41/49) |  |  |
| Kleborate Score | 49 | >4.05 | 0.44 (7/16) | 0.94 (31/33) | 0.78 (38/49) | 0.768 | 0.003 |
| Mash | 29 | <0.008 | 0.77 (7/9) | 0.9 (18/20) | 0.86<br>(25/29) | 0.900 | <0.001 |
| Jaccard | 29 | <0.318 | 0.77 (7/9) | 0.95 (19/20) | 0.90<br>(26/29) | 0.919 | <0.001 |

\*Exact test about the odds-ratio from logistic regression model
#Area under the ROC Curve
\*\*derived from classification and regression tree (CART) prediction model
\*\*\*Assessment at a potentially clinically relevant cut-point
<sup>o</sup>Computed using cut-point values

### Assessment of cKp strains that possessed all five of the biomarkers *iucA*, *iroB*, *peg-344*, *rmpA*, and *rmpA2*

Although imperfect, the presence of all five of the biomarkers *iucA*, *iroB*, *peg-344*, *rmpA*, and *rmpA2* was 94% accurate for predicting a strain that exhibited the hvKp phenotype. However, two strains, AKp14 and AKp35, both of which possessed all 5 biomarkers, exhibited a cKp phenotype as defined by the murine infection model (Supplementry Table S1). Further, both strains possessed a nearly complete hvKp-specific virulence plasmid based on Mash/Jaccard distances (Figure 2B and 2C) with the exception of an approximately 15 kb region of pLVPK missing in AKp35 as well as 18 other cKp isolates (Figure 5). Besides the pLVPK plasmid, the entire gene content of AKp14 (ST23) was compared to those of all ST23 with a confirmed hvKp phenotype in this study (AKp4, AKp10, AKp25, AKp27 and RCA1) (Supplementary Table S3). This revealed that 4,862 orthologous genes were shared by all ST23 isolates but that AKp14 uniquely carried 264 genes (due to a large insertion in the chromosome, the carriage of a prophage, and the acquisition of 3 plasmids of 45.4, 35 and 6.5 kb), and uniquely lacked 3 genes annotated as a putative ArsR family transcriptional regulator, DdpA dipeptide-binding protein, and alkanesulfonate ABC transporter permease. For strain AKp35 (ST412), no other ST412 isolate with available virulence data was available for comparison. Instead, the gene content of AKp35 was compared to all 16 strains (various ST) with confirmed hvKp phenotype in this study. While 4,203 orthologous genes were shared by all isolates, 98 genes were uniquely found in AKp35 and, importantly, 22 genes were uniquely missing in this isolate (Supplementary Table S3). Thirteen of these 22 genes were missing because of a large chromosomal deletion and belonged to a functional cluster coding for the cytochrome c maturation system (*ccmA-H*). Finally, to gain further insight into this phenotypic and genotypic virulence disparity a growth/survival analysis was performed for all 49 strains from this study in human ascites ex vivo. This medium was chosen because it mimics the in vivo growth environment and possesses complement and likely antimicrobial peptide mediated antimicrobial activity.

**Figure 5.**
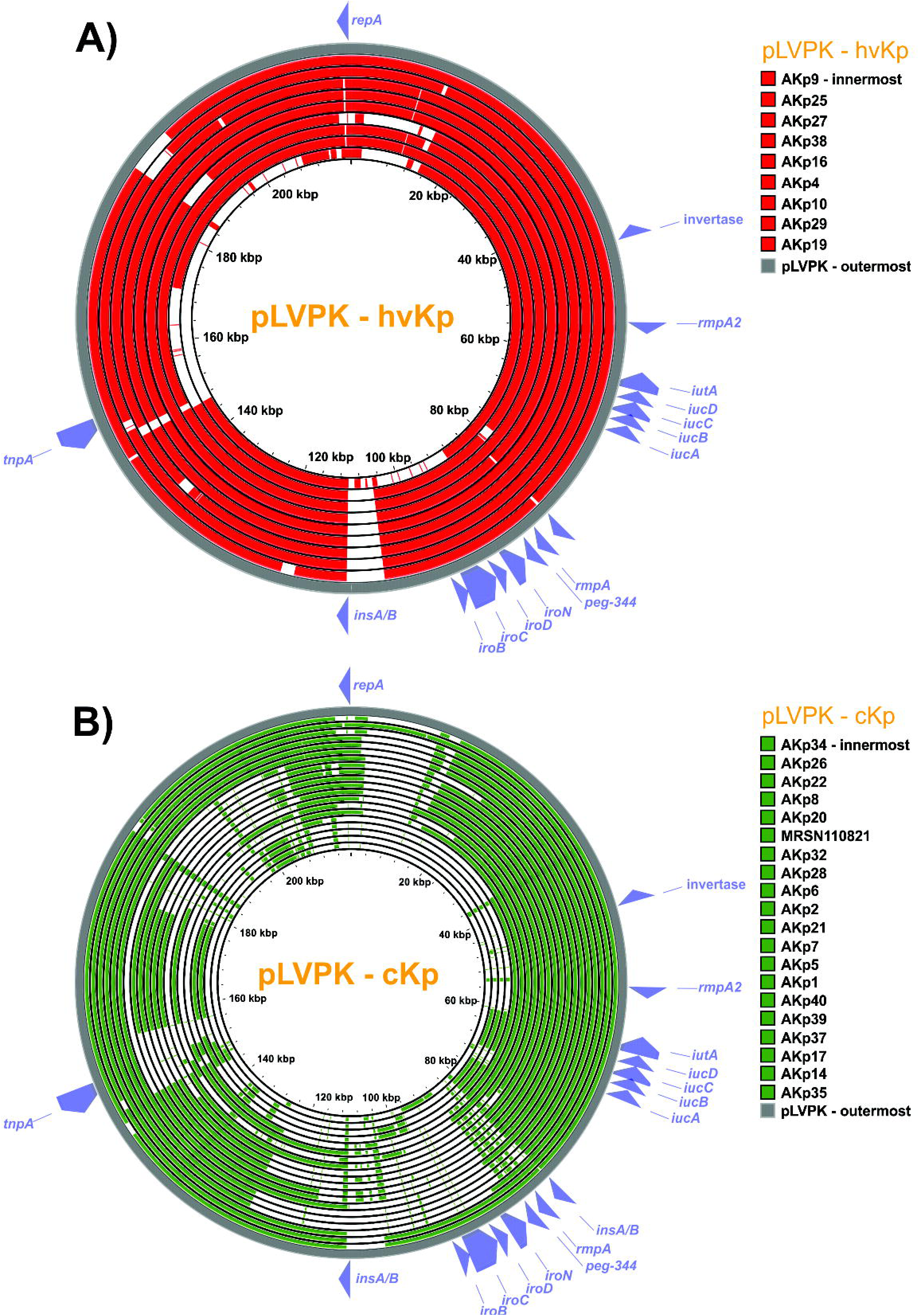
Proksee alignment to pLVPK. Twenty-nine strains with a closed plasmid harboring any virulence biomarker were compared to the canonical pLVPK. Alignment of the pLVPK sequence to all closed pLVPK-like plasmid sequences from **A)** hvKp (n=9) and **B)** cKp isolates (n=20) were produced with Proksee. Virulence and landmark genes from pLVPK are labeled in each panel. The five virulence biomarkers of interest are concentrated on a roughly 30kb region of pLVPK from *iucA* to *iroB*. Concentric circles are sorted by alignment fraction to pLVPK sequence with the outermost ring representing the isolate with the highest alignment fraction. The average alignment fraction of pLVPK-like plasmid to pLVPK is higher in hvKp isolates compared to cKp. Alignment covering the virulence biomarkers is more extensive in hvKp isolates. Most cKp isolates lack or have non-functioning *iroB, peg-344,* and *rmpA* while possessing a *rmpA2* and *iucA*.

Interestingly, by 24 hours 48/49 strains were able to grow/survive in ascites. But AKp14 was unable to do so (Supplementary Figure 1). These data support that AKp14 has a biofitness deficiency that is independent of the possession of biomarkers *iucA*, *iroB*, *peg-344*, *rmpA*, and *rmpA2* and a nearly complete hvKp-specific virulence plasmid.

## Discussion

The goal of this report was to test the hypothesis that some combination of pre-selected genotypic and phenotypic factors could differentiate hvKp and cKp strains that have acquired varying degrees of antimicrobial resistance. The effect of resistance on virulence is unresolved and may depend on the strain, the number and nature of resistance genes, and whether resistance genes on mobile elements are linked with virulence genes (32, 33). However, in some cases antimicrobial resistance may decrease virulence (34–37). Further, and more importantly, there are numerous examples of MDR– and XDR-cKp strains acquiring only a portion of the hvKp-specific virulence plasmid with the resultant pathotype designation of hvKp in the absence of their virulence potential assessed in valid infection models. Therefore, taken together, we felt it was important to assess a variety of variables that held the potential for accurately identifying the hypervirulent phenotype in this strain population. Data from this study support that the presence of all five biomarkers (*iucA*, *iroB*, *peg-344*, *rmpA*, and *rmpA2*) was most accurate in both the logistic regression model (AUC 0.962) and the CART model (94% accuracy) (Table 2). In support of this concept, Jaccard and Mash distances in comparison with the canonical pLVPK were the second-most accurate factors for predicting the hvKp strain cohort with an AUC of 0.919 and 0.90 respectively in the logistic regression model and accuracies of 90% and 86% in the CART model using the defined cut-points (Table 2).

However, it is important to note that all the strains studied in this report had acquired antimicrobial resistance beyond the ubiquitous ampicillin resistance in *K. pneumoniae*. This comparison may not be valid for strains that have not acquired additional antimicrobial resistance. These data are consistent with our earlier study in which 94% of the strains in the hvKp-rich cohort possessed all 5 biomarkers, which likely reflects the presence of a complete hvKp virulence plasmid. Notably, these strains were identified by clinical syndrome irrespective of their antimicrobial susceptibility profile (4). Interestingly, only 0.9% of a 3,123 MDR *K. pneumoniae* global strain set (27) carried all 5 biomarkers, which we find here to be most predictive of hypervirulence.

The reasonable accuracy of differentiating hvKp from cKp strains using only the five biomarkers *iucA*, *iroB*, *peg-344*, *rmpA*, and *rmpA2* was particularly encouraging. Assays for these markers could be readily developed and validated for use by clinical microbiology laboratories. It is important to note that *iutA* and *iroE* can be present on the chromosome and should not be used to differentiate hvKp from cKp (1). Phenotypic assays, such as those assessed in this study, are more challenging to implement outside of the research setting. Likewise, a whole genome genetic analysis could be equally or more insightful, but presently is best suited for research purposes and is not yet ready for use in the clinical microbiology laboratory.

The findings from this report hold promise for clinical utility. The ability to differentiate hvKp from cKp is needed for optimal clinical management. For example, hvKp infection requires vigilance for endophthalmitis and assessment for occult abscesses that could require source control. Further, the sites of infection due to hvKp could dictate modifications of the antimicrobial regimen to optimize tissue concentrations (e.g. prostate, central nervous system (CNS)) and may affect the duration of therapy. This is even more important for antimicrobial resistant hvKp (e.g. CR-hvKp) due to more limited treatment options. Although the classical clinical syndrome of an otherwise healthy individual from the community presenting with hepatic abscess, multiple sites of infection, or unusual sites of infection for cKp (e.g. endophthalmitis, CNS infection, or necrotizing fasciitis) is highly suggestive of hvKp, an increasing number of non-classical presentations, and healthcare-associated infections are being reported (38–40). This dictates the need for an accurate clinical microbiology laboratory-based test to identify hvKp. Although some laboratories have used the string test as a phenotype to identify hvKp, it lacks optimal sensitivity or specificity (4, 41, 42). The genotypic markers described in this study could form the basis of such a test, but these data demonstrate the need for all 5 biomarkers to be included (43). Although the presence of all 5 biomarkers was most accurate, using a cut-point of 4 markers versus 5 increases the sensitivity to 100% (versus 94%), albeit this comes at the cost of decreased specificity (76% versus 94%) and accuracy (84% versus 92%). Nonetheless for the clinical perspective, over-diagnosing hvKp infection could be considered more pragmatic since the consequences of missing such an infection may be significant (e.g., loss of vision).

Putative loss-of-function mutations were observed in some biomarkers and, in particular, *rmpA* and *rmpA2* due to frameshift-causing mutations in poly(G) tracks, as previously described (44, 45). Whether these presumed truncated alleles retain functionality remains to be established but, in this study, their occurrence was identical in hvKp and cKp populations. Likewise, while it cannot be excluded that putative loss-of-function mutations seen in *iuc3* plays a role in the low virulence of these strains, the remaining 19 strains with wild type *iuc* and a biomarker count ≤ 2 also exhibited low virulence in the murine model. However similar to the concept of which marker number cut-point should be utilized, even if a strain has a mutation in one of these genes that results in loss-of-function and perhaps decreased virulence, it is safer to overcall a strain as being hvKp.

Although the accuracy of predicting a strain to be either hvKp or cKp strain was 94% based on the possession of all 5 biomarkers, it was imperfect. This is due to the complexity of pathogenesis, which is a multifactorial process. The possession of defined virulence factors requisite for the hypervirulent phenotype does not necessarily reflect its overall pathogenic potential. It may be missing undefined virulence traits that were not measured, produce less surface polysaccharides which may facilitate the acquisition of resistance genes (46), or have decreased biofitness due to a variety of other factors such as antimicrobial resistance, increased sensitivity to antimicrobial factors, or other occult metabolic changes. This proved to be the case for two of the strains in our study (AKp14 and AKp35). While it remains to be functionally established, it is possible that AKp35 lost some of its virulence capabilities because of the deletion of the cytochrome c maturation system (*ccmA-H*), a chromosomal cluster ubiquitous in Gram-negative bacteria and previously linked to growth in low iron, intracellular infection, and virulence of *L. pneumophila* (47). By contrast, no major chromosomal deletions were observed in AKp14. Instead, this ST23 isolate uniquely acquired mobile elements (including 3 plasmids) accruing to a total of 264 (+5%) additional auxiliary genes. While genome reduction is a known trait associated with bacterial pathogens (48) it remains to be established whether the accumulation of plasmids in AKp14 could result in decreased biofitness.

Reinforcing this hypothesis, this strain demonstrated significantly decreased growth/survival in human ascites compared to the other 48 strains assessed in this report (Supplementary Figure S1). It is important to note that the ability to grow/survive in ascites does not equate to a hypervirulent phenotype since 32/33 cKp strains were able to do so; the hypervirulent phenotype also requires the possession of hvKp-specific virulence factors.

Conversely, the presence of some, but not all, of the established hvKp-specific virulence factor(s) does not necessarily confer the hypervirulent phenotype. Quantitative siderophore production was reasonably accurate for differentiating the hvKp-rich and cKp-rich cohorts in our initial study in which strains were identified by clinical syndrome irrespective of their antimicrobial susceptibility profile (4), but all 5 biomarkers were present in 94% of those strains. However, in this study, many strains designated as cKp possessed *iucA* and produced high levels of siderophores but possessed an incomplete repertoire of additional virulence factors and perhaps had decreased biofitness. So, the high level of siderophore production in the absence of a full armamentarium of hvKp-specific virulence factors was not sufficient to confer a hypervirulent phenotype. Hopefully future studies can begin to delineate the full repertoire of virulence factors and the genomic markers/changes that could predict an effect on biofitness, which would enable a more refined prediction of pathotypes based solely on genomics.

The Kleborate virulence score is commonly used to assess a strain’s virulence. The score ranges from 0-5 and is based on the presence of yersiniabactin (1 point), colibactin (1 point), and aerobactin (3 points) (18). However, this metric performed less well than the presence of all five biomarkers *iucA*, *iroB*, *peg-344*, *rmpA*, and *rmpA2* for differentiating the hvKp and cKp strain cohorts in this study. Using a cut-off of >4.05 derived from the CART model, the Kleborate score’s sensitivity was 44% (7/16), its specificity was 94% (31/33), and its accuracy was 78% (38/49). This may be due to the weight given to aerobactin in the Kleborate score and the fact that not all hvKp strains possess yersiniabactin or colibactin and some cKp strains possess these genes. Therefore, the Kleborate virulence score is not optimal for predicting the hypervirulent pathotype.

With the emergence of MDR and extensively drug resistant (XDR) *K. pneumoniae*, including carbapenem-resistant (CR)-Kp, an increasing number of studies have assessed for the presence of MDR-hvKp, XDR-hvKp and CR-hvKp from various strain collections (49). Such strains would represent the ultimate “superbug”, possessing both a hypervirulent pathotype and antimicrobial resistance. Many of these studies have only assessed for genotypic markers and not uncommonly only some, but not all, of the five biomarkers markers *iucA*, *iroB*, *peg-344*, *rmpA*, *rmpA2* that are evaluated in this report (5–7, 12). Although imperfect, as discussed due to potential biofitness issues, assessing for the presence of all five of the biomarkers *iucA*, *iroB*, *peg-344*, *rmpA*, and *rmpA2* will improve pathotype designation, as opposed to assessing for a more limited number of markers (43). In our study, all 24 strains that possessed only *iucA* and/or *rmpA/A2* were designated as cKp (Supplementary Table S1). Although, less pragmatic, ascites growth curves further enhance accuracy and murine infection models remain the most accurate means to identify hvKp strains (4, 50). Of note, although commonly used, the *Galleria mellonella* infection model is unable to differentiate hvKp from cKp (51).

This study has several limitations. First, these data assessed strains of *K. pneumoniae* that have acquired varying degrees of antimicrobial resistance. It is unclear whether these findings can be extrapolated to antimicrobial susceptible strains. Since antimicrobial resistance may affect biofitness, which in turn could affect a strain’s overall pathogenic potential, it is possible that antimicrobial susceptible strains may not require the possession of all 5 biomarkers to maintain a hypervirulent phenotype. Second, although we were only able to calculate Mash/Jaccard distances for 29 strains, these isolates did capture observed plasmid diversity. Although these genomic markers hold promise, calculation requires accurate plasmid construction, which cannot be achieved with Illumina sequencing alone. Lastly, the biomarkers chosen for this study were based on early data that demonstrated their utility for predicting pathotypes. However additional or alternative genotypic or phenotypic markers, both known and yet to be identified, may prove to be superior in future studies. The search for the optimal pragmatic test for resolving *K. pneumoniae* pathotypes remains a fluid process.

## Acknowledgments

This work was supported by NIH R21 AI123558-01 and 1R21AI141826-01A1 (Dr. Russo) and the Department of Veterans Affairs VA Merit Review (I01 BX004677-01) (Dr. Russo). The funders had no role in the decision to publish or the preparation of this manuscript. This study was also partially funded by the U.S. Army Medical command and the Defense Medical Research and Development Program. The authors are thankful to all the staff of the MRSN. The manuscript has been reviewed by the Walter Reed Army Institute of Research and there is no objection to its presentation. The opinions or assertions contained herein are the private views of the authors and are not to be construed as official or reflecting the views of the Department of the Army or the Department of Defense. CDC Disclaimer: The findings and conclusions in this presentation are those solely of the CDC author(s) and do not necessarily represent the views of the Centers for Disease Control and Prevention/Agency for Toxic Substances and Disease Registry. They would like to thank and acknowledge the Clinical and Environmental Microbiology Branch staff in the CDC Division of Healthcare Quality Promotion who supported this effort.

## Ethics approval and consent to participate

A subset of the isolates was collected as part of the public health surveillance activities of the MRSN, as determined by the WRAIR branch director and Human Subjects Protection Branch (HSPB) which granted ethical approval. Informed patient consent was waived as samples were taken under a hospital surveillance framework for routine sampling. The research conformed to the principles of the Helsinki Declaration. A subset of the isolates was collected under CDC surveillance and other public health activities.

## Availability of data and materials

Both genomic assemblies and raw sequencing data of the isolates analyzed in this study are publicly available in NCBI database under the Bio Project numbers PRJNA981468 and PRJNA994967.

