## Supplementary Fig. S1 for "Differentiation of hypervirulent and classical *Klebsiella pneumoniae* with acquired drug resistance"

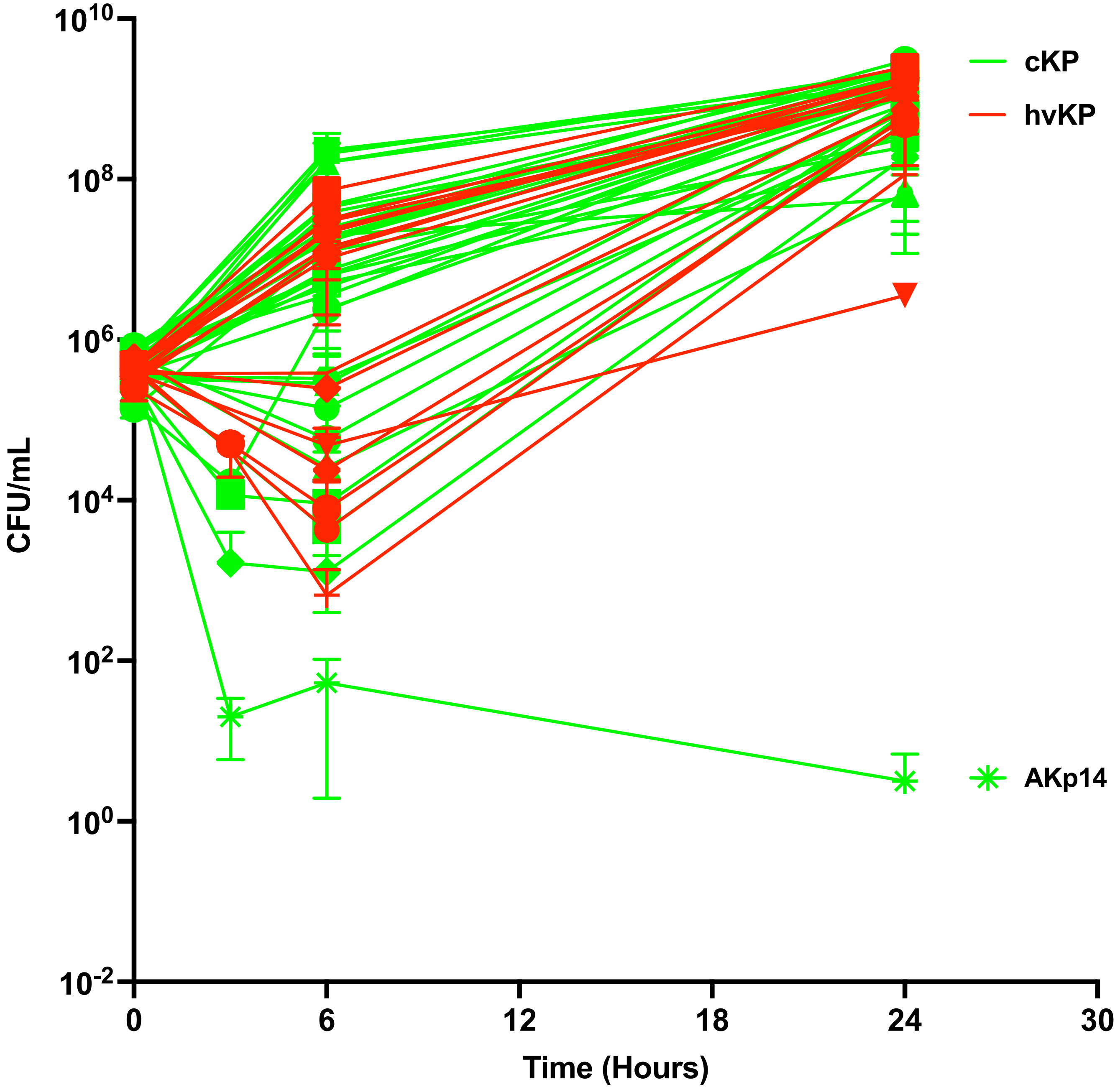


**Supplementary Figure 1**
Forty-nine strains comprising the hvKp (n=16) and cKp (n=33) cohorts underwent assessment for growth/survival in human ascites via enumeration of colony forming units. A minimum of two biologic with one-three technical repeats was performed for each strain.
